# A connectional hub in the rostral anterior cingulate cortex links areas of emotion and cognitive control

**DOI:** 10.1101/492157

**Authors:** Wei Tang, Saad Jbabdi, Ziyi Zhu, Michiel Cottaar, Giorgia Grisot, Julia Lehman, Anastasia Yendiki, Suzanne N. Haber

## Abstract

We investigated afferent inputs from all areas in the frontal cortex (FC) to different subregions in the rostral anterior cingulate cortex (rACC). Using retrograde tracing in macaque monkeys, we quantified projection strength by counting retrogradely labeled cells in each FC area. The projection from different FC regions varied across injection sites in strength, following different spatial patterns. Importantly, a site at the rostral end of the cingulate sulcus stood out as having strong inputs from many areas in diverse FC regions. Moreover, it was at the integrative conjunction of three projection trends across sites. This site marks a connectional hub inside the rACC that integrates FC inputs across functional modalities. Tractography with monkey diffusion magnetic resonance imaging (dMRI) located a similar hub region comparable to the tracing result. Applying the same tractography method to human dMRI data, we demonstrated that a similar hub can be located in the human rACC.

## Introduction

The anterior cingulate cortex (ACC) is composed of multiple regions that support a wide range of functions (emotion, motivation, higher cognition, and motor control), and thus, is in a position to use value-related information to help regulate flexibility, adaptation and top-down control (Etkin, Buchel, & Gross, 2015; Kolling et al., 2016; Shenhav, Cohen, & Botvinick, 2016). This functionally heterogeneous region is anatomically divided into the subgenual ACC (sACC), the rostral ACC (rACC), and the dorsal ACC (dACC) (Morecraft et al., 2012; Morecraft & Tanji, 2009; Ongur & Price, 2000). The sACC is connected to the motivation network consisting of the orbitofrontal cortex (OFC) and the amygdala. It is involved in visceral and emotional functions, an important mediator of motivation, and is critical for determining value (Camille, Griffiths, Vo, Fellows, & Kable, 2011; Jocham, Hunt, Near, & Behrens, 2012; Kolling et al., 2016). The rACC is tightly linked with both the sACC and the dorsolateral and ventrolateral prefrontal cortex (dlPFC and vlPFC), and is associated with cognitive control and choice of action (Jiang, Beck, Heller, & Egner, 2015; Kolling, Scholl, Chekroud, Trier, & Rushworth, 2018). Caudally, the dACC is connected with the action network consisting of motor control areas, including frontal eye fields (FEF) and premotor areas (Morecraft et al., 2012; Ongur & Price, 2000). The dACC is associated with motor planning and action execution (Caruana et al., 2018; Picard & Strick, 1996). Importantly, the connections of these divisions are a continuum and there are no specific borders between the ACC subdivisions (Morecraft et al., 2012).

The rACC sits at the connectional intersection of the motivation and action control networks. In regard to the ACC functions, the rACC is in an important position in the transition from valuation to choice to action. An important question is how the transitions occur within the rACC. One possibility is that information processing changes sequentially across subregions within the rACC, from valuation in regions close to the sACC, to cognition at the center of the rACC, and then to action in regions close to the dACC (Kable & Glimcher, 2009; Rangel & Hare, 2010; Shadlen, Kiani, Hanks, & Churchland, 2008). Alternatively, different functional processing may be integrated at a hub (Cisek, 2012; Hunt, Woolrich, Rushworth, & Behrens, 2013; Kolling, Behrens, Mars, & Rushworth, 2012; Lee, Shimojo, & O’Doherty, 2014; Rushworth, Kolling, Sallet, & Mars, 2012). Network science defines a hub as a node of a network that has a significantly larger number of links compared to other nodes in the network. In brain network analyses, the hub regions that have disproportionally numerous connections to other regions are considered central for integrating information from functionally diverse regions (van den Heuvel & Sporns, 2013). In this regard, the entire rACC can be considered a hub (Buckner et al., 2009). However, as the rACC is a large area, we sought to determine whether there is a specific region within it that is uniquely positioned to integrate signals across several functional domains.

To determine whether a hub region exists in the rACC, we systematically placed tracer injections along the rACC in nonhuman primates (NHP), and quantified the input strength and patterns from the frontal cortex (FC) to each injection site. As expected, projections from the vmPFC were concentrated in the ventral rACC and those from motor control areas were found more dorsally and caudally. Projections from the vlPFC, the dlPFC, the dmPFC and the OFC varied across sites. However, one site showed uniquely numerous and diverse connections with the FC, suggesting a hub region at this site. To test whether a similar connectivity pattern and the location of a hub could be identified in the human rACC using dMRI, we first investigated the accuracy of dMRI in detecting connectivity convergence using high resolution dMRI in NHPs. We seeded each frontal area and, consistent with the tracing results, probabilistic streamlines converged in a similar location as the hub within the rACC. We then seeded each frontal area in a dMRI dataset from the Human Connectome Project (HCP). As in the NHP results, the streamlines converged in a similar rACC location.

## Materials and Methods

### Overview

Bidirectional tracer injections were placed systematically throughout the rACC. FC was divided based on cytoarchitectonics into areas (Pandya & Seltzer, 1982; Paxinos, Huang, & Toga, 2000; Preuss & Goldman-Rakic, 1991; Vogt, 1993a; Brent A. Vogt, 2009) and retrogradely labeled cells were quantified using StereoInvestigator. To normalize for comparison across cases (variability in uptake and transport), we calculated the percentage of total labeled cells that projected from each area to each injection site. These percent scores were independent from the size of each area. The areas were further pooled into major FC regions according to their associated functions. Using the percent scores as the measurement for input strength, projection gradient across injection sites was analyzed for each FC region. Guided by the tracing results, probabilistic tractography was conducted on the NHP and human dMRI. Cortical areas were used as seed masks and the rACC as the target mask. The connectivity strength was determined by the number of streamlines between each target voxel and an FC area, divided by the total number of streamlines from that area to all rACC voxels. A convergent-connectivity value was calculated for each rACC voxel as the sum of connectivity strength weighted by the number of areas with non-zero streamlines. We identified voxels with the highest convergent-connectivity value for each individual subject and compared their locations to the hub region found in tract tracing.

### Anatomical tracing experiments

All experiments were performed in accordance with the Institute of Laboratory Animal Resources Guide for the Care and Use of Laboratory Animals and approved by the University Committee on Animal Resources at University of Rochester. Animals were adult male monkeys (*Macaca mulatta* and *Macaca fascicularis*). Bidirectional tracers were injected into the rACC. Details of the surgical and histological procedures have been described previously (Haber, Kim, Mailly, & Calzavara, 2006; Safadi et al., 2018). Monkeys were first tranquilized by intramuscular injection of ketamine (10 mg/kg) and then maintained anesthetized via 1%–3% isoflurane in 100% oxygen. Temperature, heart rate, and respiration were monitored throughout the surgery. Pre-surgery structural MR images were used to locate the stereotaxic coordinates for the injection sites. Monkeys were placed in a David Kopf Instruments stereotax, a craniotomy (2–3 cm) was made over the region of interest, and small dural incisions were made at injection sites. Bidirectional tracers (40–50 μl, 10% in 0.1 mol phosphate buffer (PB), pH 7.4; Invitrogen) were pressure injected over 10 min using a 0.5 μl Hamilton syringe, separate for each case. Tracers used for the present study were Lucifer Yellow (LY), Fluororuby (FR), or Fluorescein (FS) conjugated to dextran amine (Invitrogen). After each injection, the syringe remained *in situ* for 20–30 min.

After a survival period of 12–14 days, monkeys were again deeply anesthetized and perfused with saline, followed by a 4% paraformaldehyde/1.5% sucrose solution in 0.1 mol PB, pH 7.4. Brains were postfixed overnight and cryoprotected in increasing gradients of sucrose (10, 20, and 30%; (Haber et al., 2006)). Brains were removed and shipped to the Martinos Center for Biomedical Imaging. Diffusion MRI data was collected with the brains submerged in Fomblin solution to eliminate susceptibility artifacts at air-tissue interfaces and background signal (see *dMRI data collection and analysis* for imaging protocols). After imaging, the brains were shipped back to University of Rochester Medical Center for histological processing. Serial sections of 50 μm were cut on a freezing microtome, and one series with sections 1.2 mm-apart was processed for subsequent retrograde tracing. The serial sections were processed free-floating for immunocytochemistry. Tissue was incubated in primary anti-LY (1:3000 dilution; Invitrogen), anti-FS (1:1000; Invitrogen), or anti-FR (1:1000; Invitrogen) in 10% NGS and 0.3% Triton X-100 (Sigma-Aldrich) in PB for 4 nights at 4°C. After extensive rinsing, the tissue was incubated in biotinylated secondary antibody followed by incubation with the avidin-biotin complex solution (Vectastain ABC kit, Vector Laboratories). Immunoreactivity was visualized using standard DAB procedures. Staining was intensified by incubating the tissue for 5–15 s in a solution of 0.05% DAB tetrahydrochloride, 0.025% cobalt chloride, 0.02% nickel ammonium sulfate, and 0.01% H_2_O_2_. Sections were mounted onto gel-coated slides, dehydrated, defatted in xylene, and coverslipped with Permount. In cases in which more than one tracer was injected into a single animal, adjacent sections were processed for each antibody reaction.

### Analysis: strength of inputs and defining the hub

Seven out of sixteen injection sites were selected for analysis based on the following criteria (Fig. 1): 1. Location of the injection site along the rACC; 2. lack of tracers leaking into adjacent cortical regions or into the white matter; 3. outstanding transport; and 4. low background. Two cases were in the same position, however, one particularly large, and served as a control for size. This case demonstrated that the number and strength of projection to a site was not dominated by the injection size, but rather its position. To evaluate the strength of the different FC inputs to each injection site, we divided the FC into 27 areas based on the atlas by Paxinos et al. (2000), in conjunction with detailed anatomical descriptions (Pandya & Seltzer, 1982; Preuss & Goldman-Rakic, 1991; B. A. Vogt, 2009b; Vogt, 1993b). Labeled cells were quantified throughout the prefrontal cortex (PFC) and the premotor cortex using StereoInvestigator software (MicroBrightField) as previously described (Choi, Tanimura, Vage, Yates, & Haber, 2016). To compare the input pattern across injection sites, the percent input to each site was calculated based on the number of labeled cells in each FC area projecting to a given site, divided by the total number of labeled cells across all FC areas projecting to the same site. Areas were then ordered based on their percent scores. The number of areas whose cell counts added up to 50% and 75% of total input was calculated for each site. To determine whether the inputs were distributed evenly across areas or highly concentrated in a few areas, an entropy score was calculated to compare the input pattern to a uniform distribution:

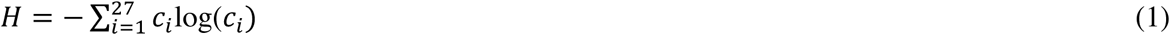
where *c_i_* is the percent cell count for the *i*-th area, and log() is the natural logarithm function (Conrad, 2004). The hub was characterized by a high number of areas contributing to 50% and 75% of total inputs. Additionally, the hub was expected to have a high entropy score that indicates evenly distributed inputs across areas.

**Figure 1.**
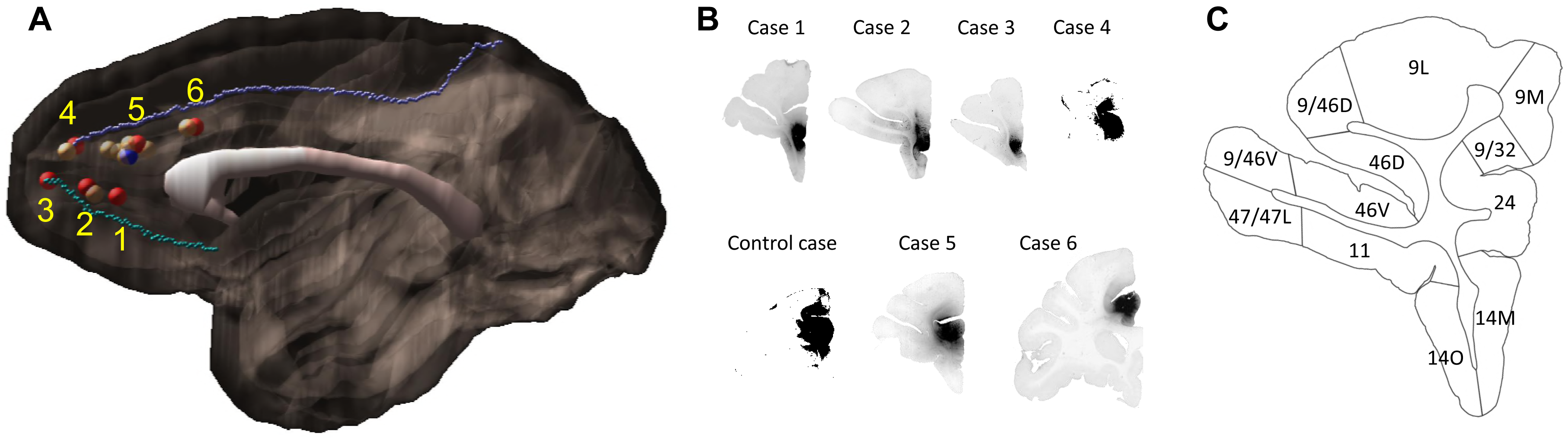
Injection sites and cortical area definition. (A) Locations of the 7 out of 16 ACC injections that were selected for stereology analysis. Six cases (red) were analyzed as the main result and the additional one (blue) as control. The numbering of cases followed the longitudinal axis of the cingulate gyrus. Cases not used for stereology due to the limitation in transport, background and location were colored beige. The cortex of the right hemisphere is shown in semi-transparent brown, the corpus callosum in white, and the cingulate sulcus and rostral sulcus in purple and cyan. (B) Photomicrographs to show the histology of each injection site. (C) Section outline of FC areas on an example slide. The borders between areas were hand-drawn following the atlas by Paxino et al. (2008).

### Definition of commonly referred regions

To determine whether inputs to each site show spatial regularity in their cortical origin, FC areas were grouped into regions commonly used in the terminology of functional studies (see e.g. (Clark, Boutros, & Mendez, 2010)). These included: frontal pole (FP, area 10), vmPFC (areas 14M, 25), OFC (areas 14O, 11, 13, OPAl, OPro), vlPFC (areas 47, 47O, 44, 45), dlPFC (areas 9, 46, 9/46), dmPFC (areas 9/32, 9M), frontal eye field (FEF, area 8) and premotor cortex (areas 6, ProM, 6/32). We note that in many functional imaging studies in human, the demarcation of vmPFC includes the medial part of Broadmann Area 10. However, area 10 in both NHPs and humans has a dorsal and a lateral part that covers the entire polar region. To maintain cytoarchitectonic consistency in spatial demarcation, we separated area 10 from vmPFC entirely and designated it as FP.

### dMRI data collection and analysis

The dMRI data collection and preprocessing was previously described in Safadi et al. (2018). The NHP dMRI data was collected from 7 animals in a small-bore 4.7T Bruker BioSpin MRI system, with gradient internal diameter of 116 mm, maximum gradient strength 480 mT/m, and birdcage volume RF coil internal diameter of 72 mm. A 3D echo-planar imaging (EPI) sequence was used for dMRI with TR = 750 ms, TE = 43 ms, δ = 15 ms, ∆ = 19ms, b_max_ = 40,000 s/mm^2^, 514 gradient directions, matrix size 96 × 96 × 112, and 0.7 mm isotropic resolution. The human dMRI data used 35 healthy subjects, publicly available as part of the MGH-USC Human Connectome Project (HCP) (Fan et al., 2016). Both the NHP and human data were preprocessed using FSL 5.0.9 (Jenkinson, Beckmann, Behrens, Woolrich, & Smith, 2012). Artifacts of head movements and distortions by eddy currents were corrected(Andersson & Sotiropoulos, 2016). A crossing fiber model (bedpostx)(Behrens, Berg, Jbabdi, Rushworth, & Woolrich, 2007) was fit to each voxel to estimate the distribution of fiber orientations, which were subsequently used in the probabilistic tractography.

Following preprocessing, probabilistic tractography (Behrens et al., 2007) was performed in each individual’s diffusion space, and the results were transformed to a template for comparison across subjects. Parcellation of the FC areas for NHP followed the same atlas used for tract tracing (Paxinos et al., 2000). Each FC area was used as a seed, while ACC areas 32 and 24 were combined and used as the target. Twenty-seven seed masks were created correspondent to the 27 FC areas used in the tracing analysis. To generate seed and target masks, areal masks from a template brain (Calabrese et al., 2015) were transformed to each individual’s diffusion space via nonlinear registration (Klein, Staring, Murphy, Viergever, & Pluim, 2010). Each mask covers 0.14 mm thickness (2 voxels) of white matter at the gray-white matter boundary (Fig. 7B). Anatomical parcellation of the human cortex was hand drawn on a surface-based template (*fsaverage* by FreeSurfer 4.5, Fig. 8A). The parcellation followed Petrides et al. (2012), which was developed to maximize architectonic correspondence between human and NHP prefrontal areas (see Discussion on the cross-species homologies). Twenty-five hand-drawn masks were transformed from the *fsaverage* space to each individual’s diffusion space with nonlinear registration provided by FreeSurfer. The FC areas were used as seeds, and areas 32 and 24 were combined and used as the target. Each mask covers 3 mm thickness (2 voxels) of white matter at the gray-white matter boundary.

FSL was used for probabilistic tractography for both NHP and human. Tractography was performed from each seed mask to the ipsilateral target mask. At each voxel, a sample was drawn from the orientation distribution of the anisotropic compartment with the closest orientation to the previously visited voxel (Behrens et al., 2007). To exclude indirect pathways through subcortical structures, the thalamus, the striatum and the amygdala were used as exclusion masks. To measure the connectivity strength between each seed mask and each target voxel, the number of streamlines arriving at the target voxel was divided by the total number of streamlines generated from the seed mask. The convergent-connectivity value was defined as the sum of areas with non-zero streamlines multiplied by the connectivity strength. The final results from all individuals were displayed on a template brain via linear registration (FA image of the template by Calabrese et al., 2015 for NHP, T1-weighted image of the MNI152 template for human).

## Results

### Overview

Injection sites were labeled 1–6 based on the spatial order of their locations (Fig. 1). Sites 1–3 were in area 32 and labeled from caudal to rostral respectively. Site 4 was in the transition zone between areas 32 and 24. Sites 5 and 6 were in area 24. Sites 4–6 were labeled from rostral to caudal. The additional large injection was in area 24 and overlapped with site 5, but, extended beyond the borders of site 5. This case served as a control for the variability of injection size. Retrogradely labeled FC cells were quantified in each case for each area. The projection strength from each cortical area was calculated based on the percent of total labeled cells. The sites differed with respect to projection strength from various FC areas to each site. The results demonstrated 3 projection patterns across sites: a decrease in the input strength to sites 1–6 from the FP and the vmPFC; an increase in the input strength to sites 1–6 from the FEF and the premotor cortex; a non-monotonic gradient centered on site 4 with the input strength from the dlPFC, the dmPFC and the vlPFC increasing to sites 1–4 and decreasing to sites 4–6. Taken together the results show that site 4 receives inputs from the greatest number of FC areas. It also contains the most regionally diverse inputs compared to the other sites. Finally, consistent with the tracing results, dMRI tractography in NHP and human showed convergent probabilistic tracts from the FC to an rACC region in close proximity to site 4.

### Projection patterns from the FC areas to the rACC

Not all FC regions project equally to the rACC. Overall, retrogradely labeled cells in cases 1–3 were primarily located in more rostral regions of the FC, while those in cases 4–6 were primarily in more caudal regions (Fig. 2). Case 1 showed high concentration of labeled cells at the rostral pole of the medial PFC. However, at more caudal levels, the concentration of labeled cells was located primarily ventrally, with few labeled cells in areas 9 and 9/32. In addition, there were clusters of labeled cells located in a caudal OFC area OPro. Similarly, in cases 2 and 3, labeled cells were concentrated rostrally in the FP and the vmPFC regions. However, some clusters with a few labeled cells were also located more dorsally in area 9L and caudally in the lateral OFC. In contrast for case 1, additional labeled cells were found in the vlPFC (areas 47L and 47O). Case 4 showed a diverse pattern to the distribution of labeled cells, with a clear increase of dense clusters of labeled cells located in dorsal regions. At the most rostral level, labeled cells were concentrated in the FP. At more caudal levels, labeled cells were extensively distributed in the dlPFC, the vlPFC, ventromedial OFC and the vmPFC. An additional cluster of labeled cells was also found in area OPro. Case 5 resulted in labeled cells concentrated ventrally, primarily in the caudal part of orbital and medial regions, although scattered clusters of labeled cells were also located rostrally in the FP and dorsally in area 9. Finally, labeled cells in case 6 were primarily distributed in caudal dorsal regions, in area 8 and the premotor cortex. Only very few labeled cells were found in ventral or rostral regions.

**Figure 2.**
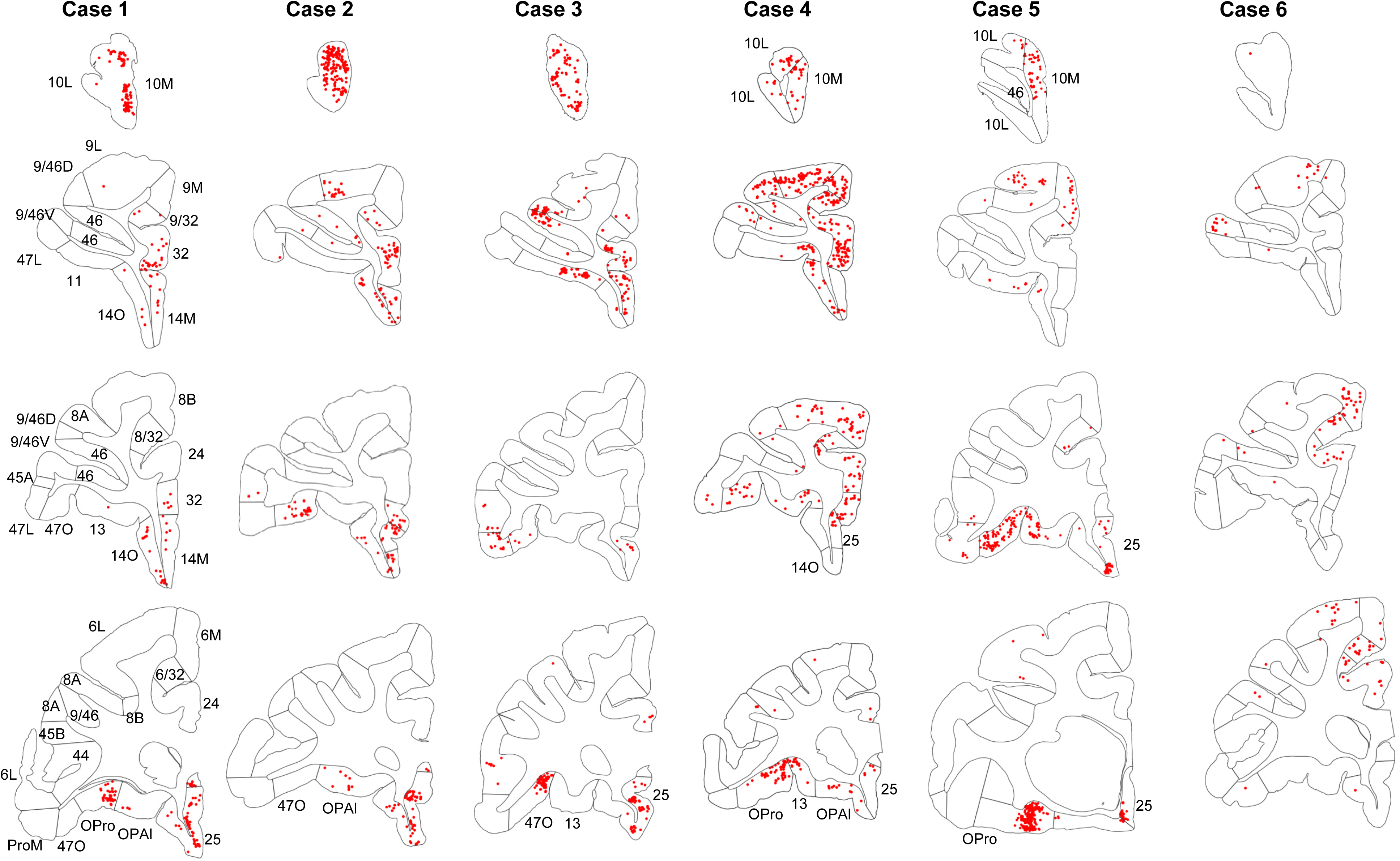
Retrogradely labeled cells (red) in cases 1–6. Each column contains rostral to caudal coronal sections from one case. Sections of the same row have matching locations along the rostro-caudal axis. FC areas are labeled in the first column. Areas not matching the parcellation in the first column were labeled additionally on the corresponding sections.

An essential property of network hubs is their high degree of connections. To measure the degree of connection of each site, we identified the number of areas that contributed to 50% and 75% of total inputs. This number varied extensively across sites (Fig. 3). At site 1, only two areas, 25 and 10M accounted for 50% of the total inputs. Inputs from areas 10L and OPro accounted for an additional 25%, for a total of 4 areas comprising 75% of all inputs to site 1. Similarly, two areas, 10L and 10M contributed 50% of total inputs to site 2, and two areas 25 and 46 accounted for an additional 25%, for a total of 4 areas comprising 75% of all inputs to site 2. At site 3, three areas 25, 46 and 47O contributed 50% of the total inputs, with three additional areas 9L, 10L and 11 (the additional 25%) for a total of 6 areas comprising 75% of all inputs to site 3. In contrast to sites 1–3, site 4 received 50% of the total inputs from five FC areas 9L, 47O, 9M, OPro and 46. Five areas, 9/32, 14O, 14M, 8B and 9/46D, contributed the additional 25%, for a total of 10 areas comprising 75% of all inputs to site 4. More consistent with sites 1–2, site 5 received 50% of its inputs from only two areas, OPro and 47O. Four areas, 13, 10M, 25 and 9L contributed the remaining 25%, for a total of 6 areas comprising 75% of all inputs to site 5. The control site was large and had a higher number of labeled cells than all the other sites (Fig. S1A). Despite that, it showed a similar distribution of major input areas to that of site 5. The control site received 50% of its inputs from two areas, 13 and 47O. Three areas, OPAl, 47L and 46 contributed the remaining 25%, for a total of 5 areas comprising 75% of all inputs (Fig. S1B). Finally, four areas, 6L, 6/32, 9/46V and 8B, contributed 50% of total inputs to site 6. Five areas, 6M, 8A, ProM and 8/32 contributed an additional 25%, for a total of 8 areas comprising 75% of all inputs to site 6. In summary, the number of areas accounting for 50% and 75% of total inputs varied across sites (Table 2). Importantly, this was not a function of the differential size of the areas (Fig. S1C). The curves generated by calculating the number of areas contributing 50% of the total inputs to each site illustrate the relatively limited number of input to sites 1–3 and 5 compared to sites 4 and 6 (Figs. 3 & 4A). The entropy measure of cell counts across areas to verify that the difference of cell distribution across sites was not due to an arbitrary percentage cutoff. Consistent with the area counts in Fig. 4A, the entropy score was the lowest at sites 1 and 2, and the highest at site 4 and site 6 (Fig. 4B).

**Figure 3.**
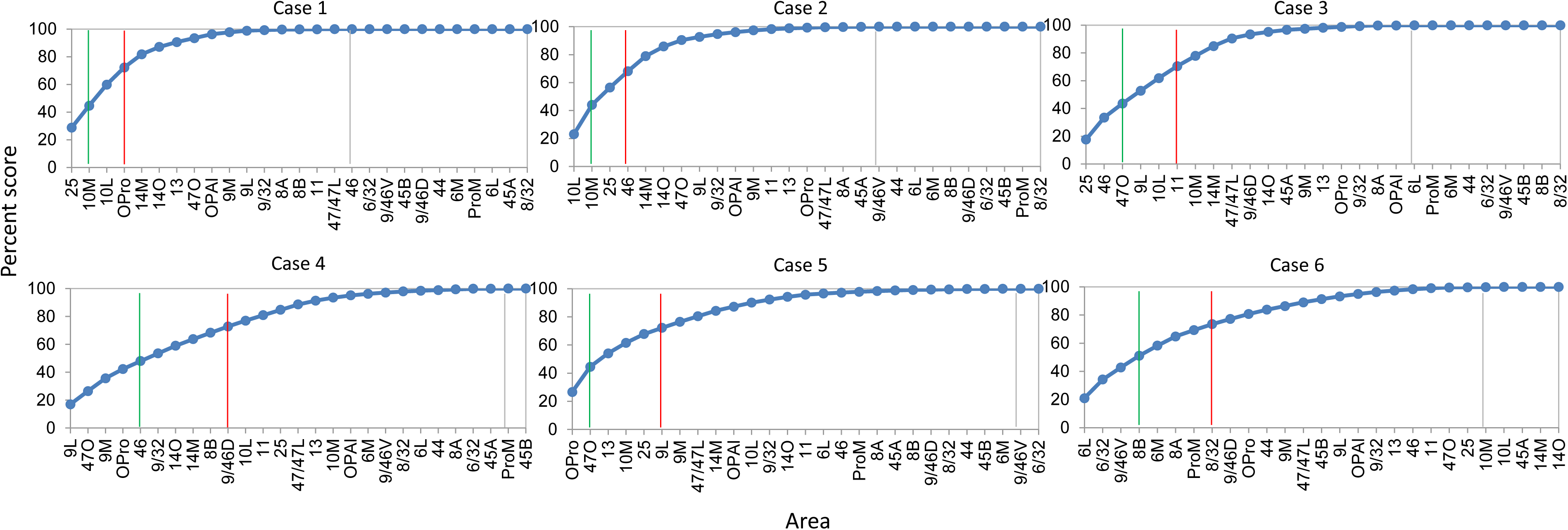
Cumulative percent cell count across areas in cases 1–6. Cutoff remarks: green line = 50%, red line = 75%, grey line = 100%. Areas after 100% are in a random order. The slope of each curve indicates deviation from a uniform distribution (steeper slope = further deviation).

**Table 1:** List of the 27 frontal areas.

| vmPFC | dmPFC | OFC | vlPFC | dIPFC | FEF | Premotor |
| --- | --- | --- | --- | --- | --- | --- |
| 10L | 9M | 11 | 47/47L | 9/46V | 8A | 6L |
| 10M | 6M | 13 | 47O | 9/46D | 8B | 6/32 |
| 14M | 9/32 | OPro | 44 | 46 | 8/32 | ProM |
| 14O |  | OPAI | 45A | 9L |  |  |
| 25 |  |  | 45B |  |  |  |

**Table 2:** Summary of the major FC inputs to each site.

| Site | # areas contributing<br>50% inputs | Regions contributing<br>50% inputs | # areas contributing<br>75% inputs | Regions contributing<br>additional 25% inputs |
| --- | --- | --- | --- | --- |
| 1 | 2 | FP, vmPFC | 4 | FP, OFC |
| 2 | 2 | FP | 4 | vmPFC, dIPFC |
| 3 | 3 | vmPFC, vlPFC, dIPFC | 6 | FP, OFC, dIPFC |
| 4 | 5 | OFC, vlPFC, dmPFC,<br>dIPFC | 10 | vmPFC, OFC, dmPFC,<br>dIPFC, FEF |
| 5 | 2 | OFC, vlPFC | 6 | FP, vmPFC, OFC, dIPFC |
| 6 | 4 | dIPFC, FEF, Premotor | 8 | FEF, Premotor |

**Figure 4.**
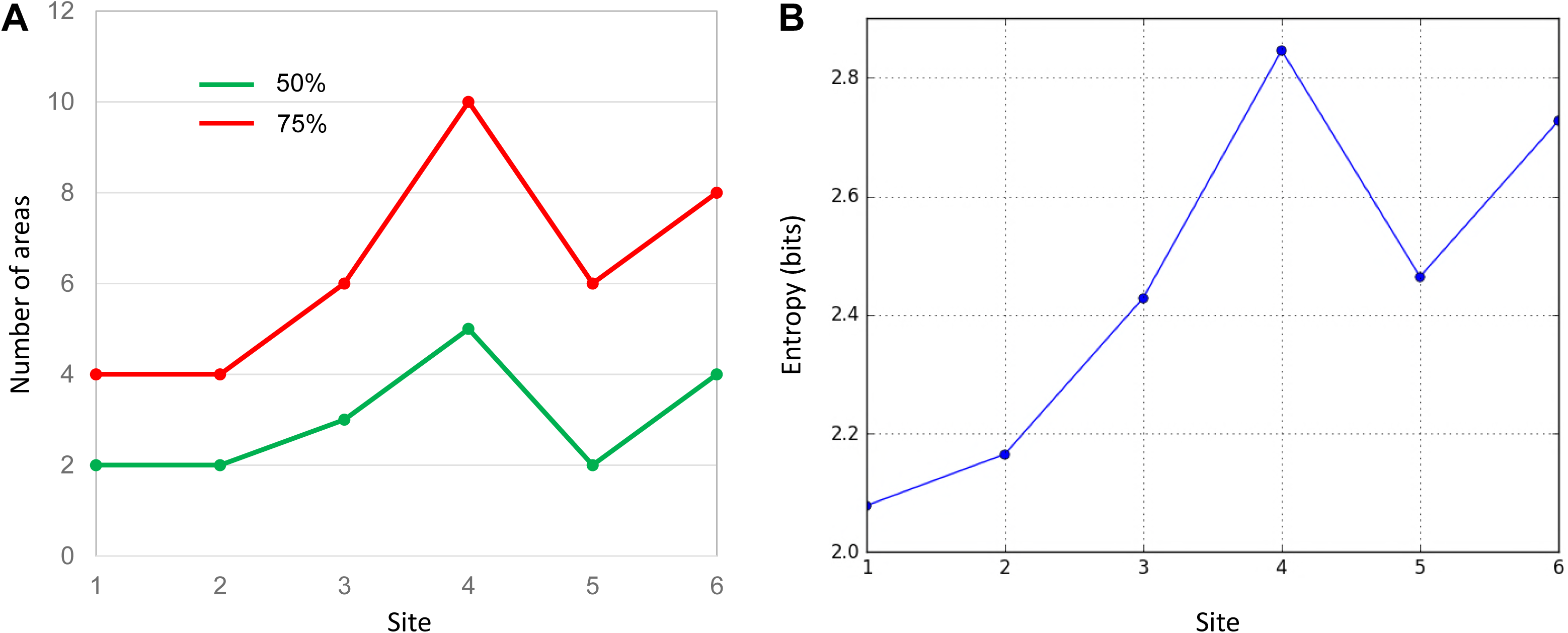
Quantitative comparison of cell distributions in cases 1–6. (A) The number of areas that contributed 50% (green) and 75% (red) of inputs in each case. The highest number was found in case 4. (B) The entropy of cell distribution in each case. Higher entropy corresponds to less deviation from the uniform distribution, i.e. more evenly distributed labeled cells across areas. The highest entropy was in case 4.

### Projection patterns from cortical regions to the rACC

FC areas were grouped into commonly referred FC regions. These included: frontal pole (FP, area 10) and vmPFC (areas 14M, 25), OFC (areas 14O, 11, 13, OPAl, OPro), vlPFC (areas 47L, 47O, 44, 45), dlPFC (areas 9, 46, 9/46), dmPFC (areas 9/32, 9M), frontal eye field-FEF (area 8), and premotor cortex (areas 6, ProM, 6/32). At each site, we identified the FC regions contributing 50% and 75% inputs (Fig. 5). FP and vmPFC contributed 50% of the total inputs to site 1. The next 25% were from FP and OFC, for a total of 3 regions. At site 2, only FP contributed 50% of the inputs. Inputs from vmPFC and dlPFC comprised the next 25% for a total of 3 regions. vmPFC, vlPFC and dlPFC contributed 50% inputs to site 3, and OFC and FP the next 25% for a total of 5 regions. OFC, vlPFC, dmPFC and dlPFC contributed 50% of the total inputs to site 4. The next 25% were from vmPFC, OFC, dlPFC and FEF, for a total of 8 regions. Inputs from OFC and vlPFC comprised 50% of the inputs to sites 5. The next 25% were from FP, vmPFC, OFC, and dlPFC, for a total of 5 regions contributing to 75% of all the inputs to site 5. Finally, at site 6, both the 50% and 75% inputs were contributed by 3 regions: premotor cortex, FEF and dlPFC. In summary, sites 1, 2 and 5 had the most limited regional input, and site 4 had the most diverse regional input.

**Figure 5.**
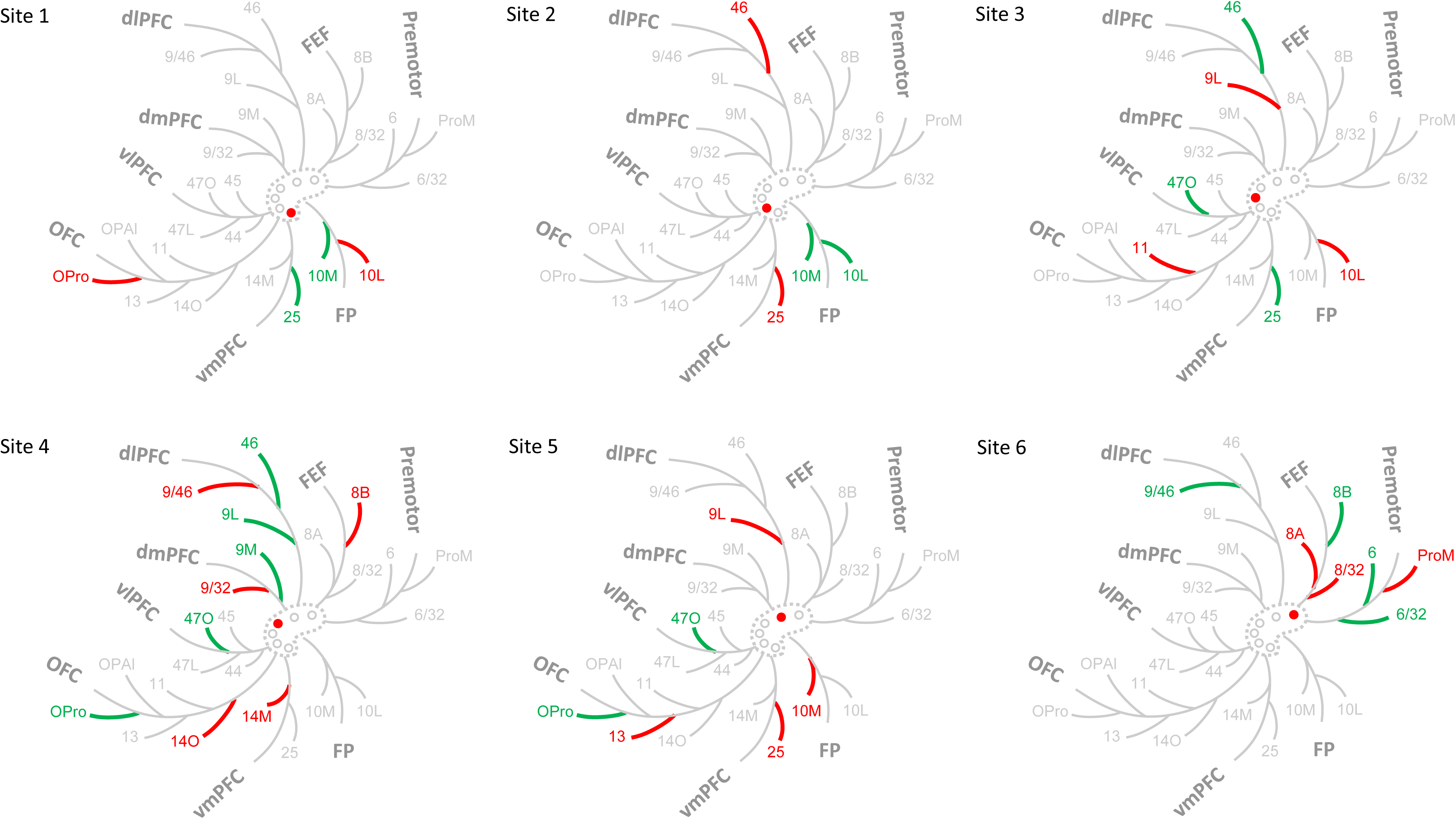
Schematic illustration of the FC regions with strong projections in each case. The dashed contour represents the ACC and the circles injections of cases 1–6. The filled circle marks the case being shown. Colored branches represent brain areas that accounted for 75% cell counts in each case (green = adds up to 50%; green + red = adds up to 75%). The most extensive FC regions with strong input was found in case 4.

### Projection trends from cortical regions to the rACC

Using the mean percent score for the projection strength from each cortical region to each site, we identified three modes of cortical projection patterns to the rACC (Fig. 6). There were two monotonic trends across all sites, one related to the FP and vmPFC projections, the other the premotor and FEF projections. The third mode was a nonmonotonic gradient with a single peak at site 4, related to the dlPFC, vlPFC, and dmPFC projections. In the first trend, vmPFC and FP contributed to more than 15% of inputs to sites 1 and 2, ~10% to site 3, just under 5% to sites 4 and 5, and close to 0% to site 6 (Fig. 6A). The between-site difference was statistically significant (Kruskal-Wallis *H* = 10.48, *p* < 0.02). This trend indicates a decrease in projection strength of vmPFC and FP along a gradient from site 1 to 6. In contrast, the second trend demonstrates an increase of projection strength from site 1 to 6 of inputs from premotor cortex and FEF. These regions projected strongly to site 6, but weakly to the other sites. Indeed, the average contribution of an area in these regions was less than 1% to the inputs to sites 1–3, ~2% to sites 4 and 5, and more than 5% to site 6 (Fig. 6B). The between-site difference was also statistically significant (Kruskal-Wallis *H* = 35.46, *p* < 1×10^−5^). Unlike projections from FP and vmPFC, this trend is less evenly distributed along a gradient from site 1-6, as the projections are concentrated in site 6. The final projection pattern is a single-peak nonmonotonic gradient centered at site 4. The average input strength from dlPFC, vlPFC and dmPFC increases from site 1 (~1%) to site 4 (> 5%) and then decreases from site 4 to site 6 (~2%). The change across sites is more gradual than that in the 2 monotonic trends. The between-site difference was statistically significant (Kruskal-Wallis *H* = 20.18, *p* < 0.01). Interestingly, OFC areas did not show a trend across the rACC (Fig. 6D). Rather, the OFC projection was strongest to site 5 (8%), 1 (5%), and 4 (4%) and less so to sites 2 (2%) and 6 (1%).

**Figure 6.**
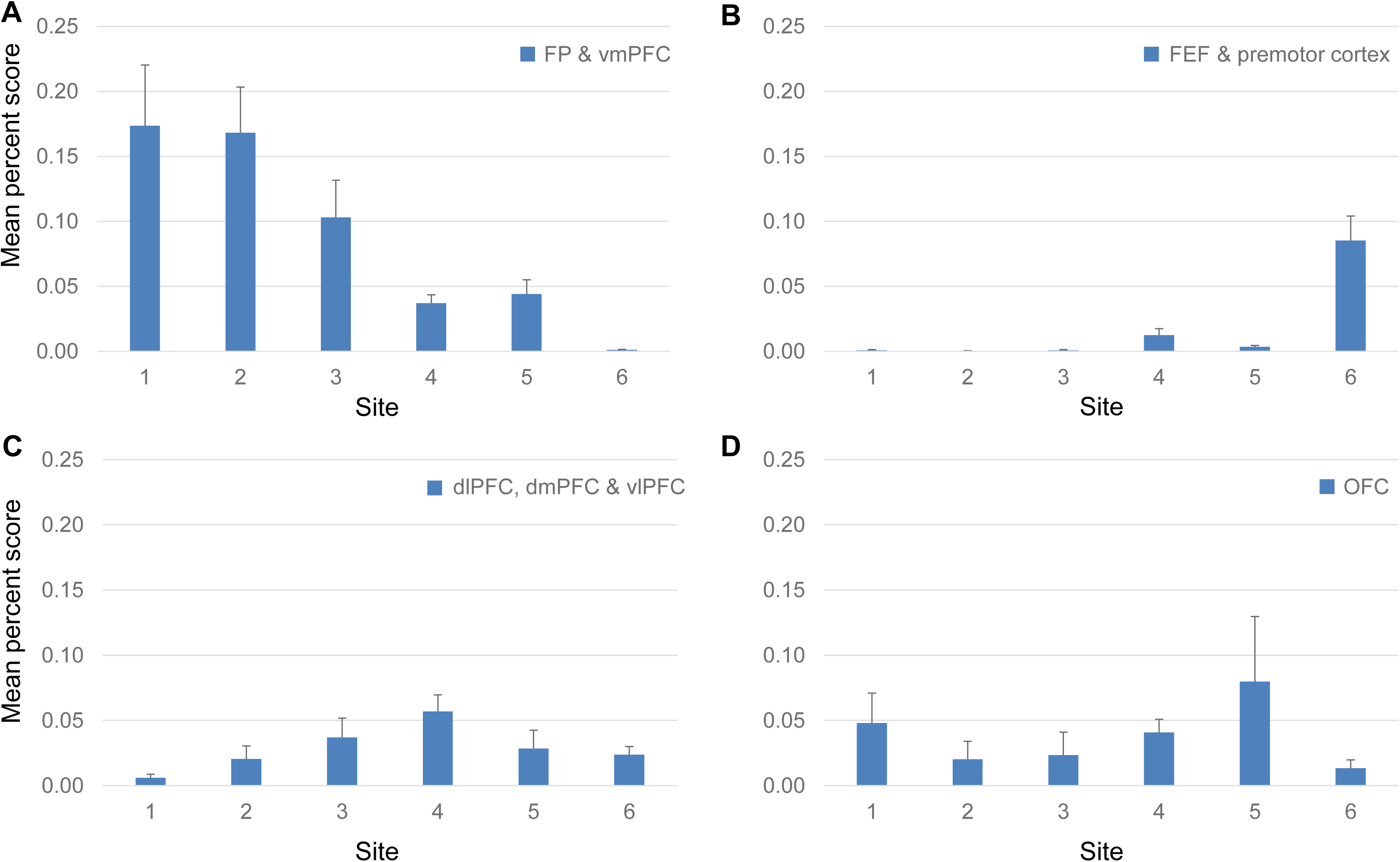
Projection strength from different FC regions at each site. Percent scores of inputs from (A) FP & vmPFC, (B) FEF & premotor cortex, (C) dlPFC, dmPFC & vlPFC and (D) OFC are shown in separate panels. In each panel, the percent scores of areas in the corresponding regions were averaged. The mean and standard error across areas are shown for each site. The mean percent score of FP & vmPFC was greater at sites 1-3 than at sites 4-6; that of FEF and premotor cortex was lower at sites 1–3 than at sites 4-6. The mean percent score of dlPFC, dmPFC and vlPFC gradually increases from site 1-4 and decreases from site 4–6. There was no consistent pattern in the OFC percent scores across sites.

### Summary

FC inputs to 6 sites in the rACC varied with respect to strength and regions of origin (Table 2). Site 1 & 2 receive the strongest (50%) inputs from 2 areas (vmPFC and FP); site 3 receives the strongest inputs from 3 areas (vmPFC, vlPFC and dlPFC); site 4 receives the strongest inputs from 5 areas (OFC, vlPFC, dmPFC and dlPFC); site 5 receives the strongest inputs from 2 areas (OFC and vlPFC); and site 6 receives the strongest inputs from 4 areas (dlPFC, FEF and premotor cortex). Importantly, site 4 stands out as having the highest number of areas that contributed 50% and 75% of inputs to it. Site 4 also had the most diverse cortical inputs. Across sites, the projection strength of different regions formed 3 spatial patterns: vmPFC and FP showed increasing projection strength from site 1 to 6; FEF and premotor cortex showed decreasing projection strength from site 1 to 6. The third pattern was a nonmonotonic gradient formed by projections from dlPFC, dmPFC and vlPFC, with a single peak centered at site 4.

### Convergent probabilistic tracts from the FC to the rACC in dMRI

The FC of NHP dMRI images was parcellated into 27 areas that corresponded to the 27 areas used in the tracing analysis (Fig. 7A). Each area was used as a seed mask. Areas 24 and 32 were combined as the target mask. Probabilistic streamlines from the different seeds terminated in partially overlapping regions in the target mask. Each streamline was a probabilistic estimation of the path that connects a seed voxel and a voxel in the ACC. As an example, Fig. 7B shows two seed masks from areas 11 and 46, and Fig. 7C illustrates the voxels where streamlines from the two seed masks terminate in the ACC in one monkey. A subgroup of the streamlines from both areas targeted the same voxels (shown in orange in Fig. 7C). We identified the location of highest convergence, i.e. the voxel receiving streamlines from the most number of seeds. A convergent-connectivity value was calculated for each voxel in the target mask, approximating the number of areas with high density of streamlines to that voxel. Consistently across 7 monkeys, the highest convergence-connectivity value found across all animals was located at the rostral edge of the cingulate sulcus (Fig. 7D). This is in a similar location as site 4 in the tracing experiments (see Fig. 1A). There was some individual variability on the dorsal-ventral axis. In 3 animals, the highest convergent-connectivity value was just dorsal to the cingulate sulcus and in 4 animals it was just ventral to the sulcus.

**Figure 7.**
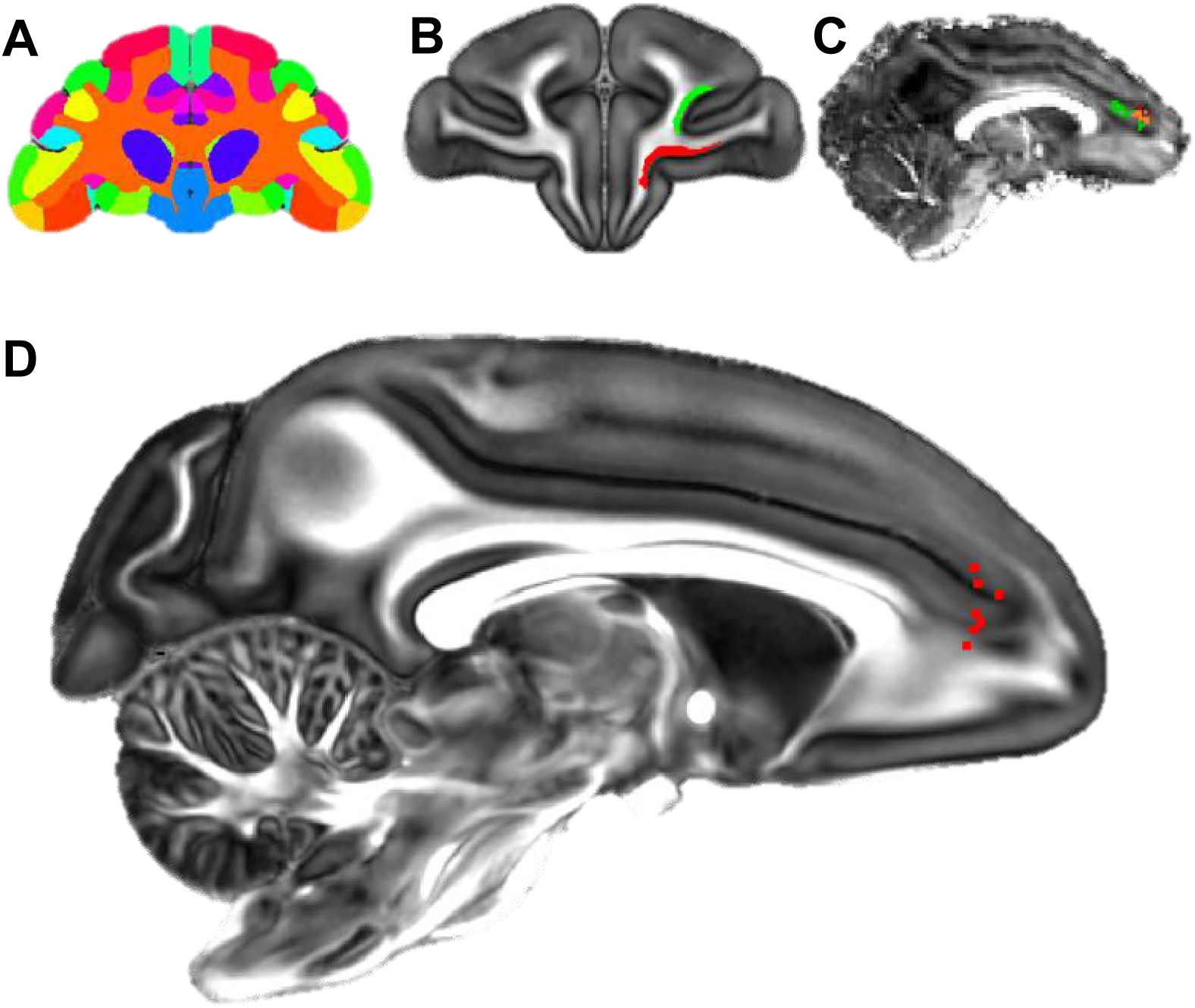
Localization of the hub region in the monkey rACC using dMRI tractography. (A) A coronal section illustrating the Paxinos atlas in the dMRI space provided by Duke University. (B) A coronal section of the atlas brain showing two example seed masks for areas 11 (red) and 46D (green). (C) A sagittal section of an individual monkey brain showing the probabilistic streamline terminals in the rACC, separately for the seeds in area 11 (red) and 46D (green). Voxels with overlapping terminals were in orange. (D) A sagittal section showing the localized hub in 7 individual monkeys. Each red dot marks the center of the hub region in one monkey. The center of the hub was defined by the voxel with the highest weighted-sum of probabilistic streamlines from all 29 seeded areas.

We applied the same tractography method to human dMRI data. The human FC was parcellated into 25 areas following Petrides and Pandya (1994). This anatomical division was developed to maximize architectonic correspondence between human and NHP frontal areas (OPro and OPAl were not clearly defined in the human parcellation by Petrides and Pandya (1994), and were thus not included in our human dMRI analysis). Similar to the NHP analysis, areas 32 and 24 were combined as the ACC target mask. Probabilistic streamlines were generated from each seed to the target. A convergent-connectivity value was calculated for each voxel in the target mask. The results demonstrated that, as in the NHP results, streamlines in each subject converged in the rACC. The highest convergent-connectivity value was consistently located in the rostral part of the rACC for all subjects (Fig. 8). The geometric center of the individual results was at the genu of the cingulate gyrus. This region was spatially approximate to site 4 in the NHP tracing study. Despite the morphological difference of cingulate sulcus between human and NHP, the geometric center of the results in Fig. 8 was at the rostral edge of the human cingulate sulcus, based on the Mai human atlas (Mai, Paxinos, & Voss, 2008). Interestingly, as in the NHP data, there was little individual variance in the rostrocaudal location of the convergence. However, similar to the NHP results, there was some individual variation in the dorsal-ventral axis. Importantly, as seen in NHP data, about half of the subjects had the voxel of highest convergence above the genu, and the other half below the genu.

**Figure 8.**
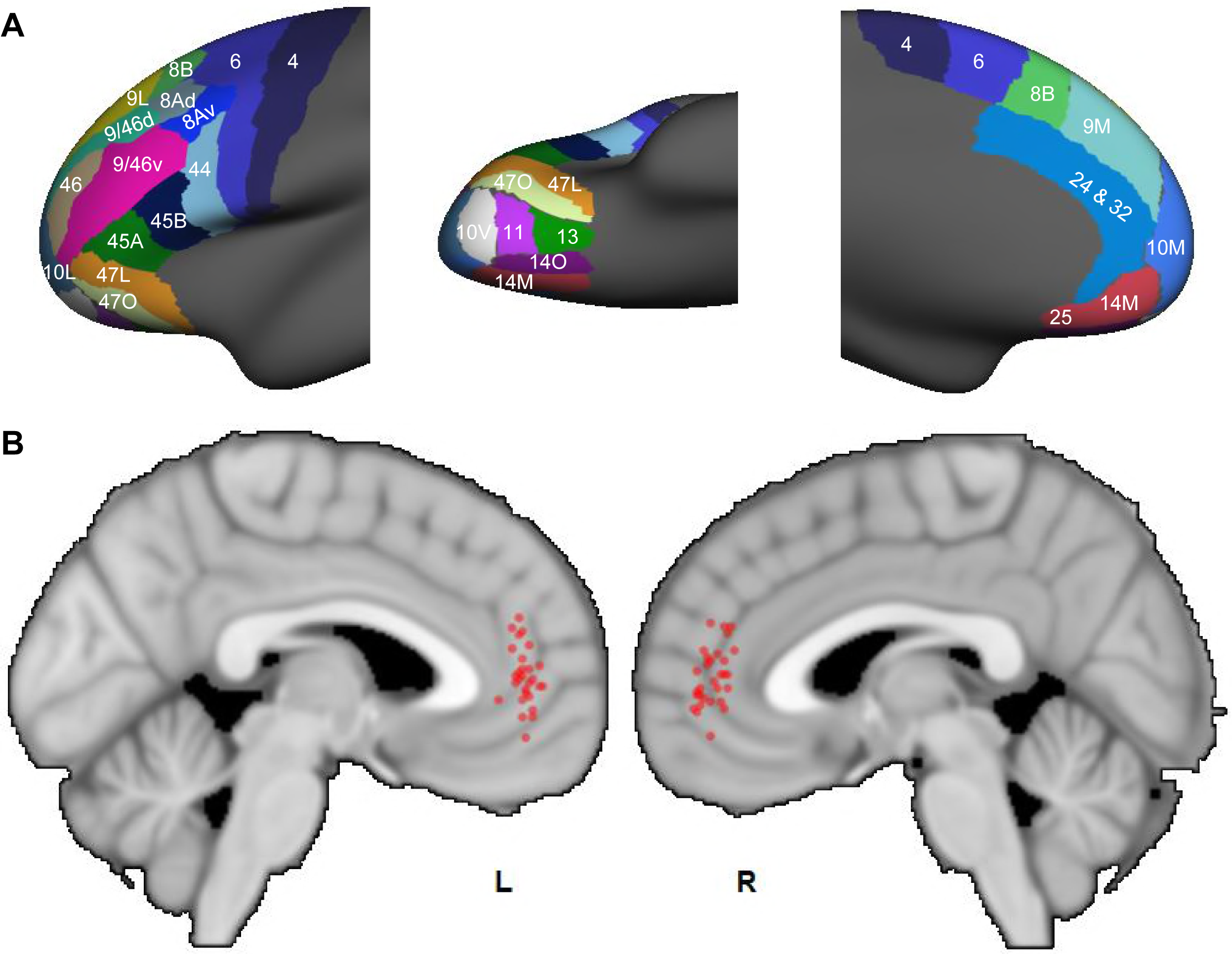
Probabilistic streamlines converging in the rACC in human dMRI. (A) Parcellation of the FC areas on the *fsaverage* template (FreeSurfer 4.5), following Petrides et al. (2012). The FreeSurfer labels are available in the Supplementary File. (B) Sagittal sections showing the localized hub across individuals. Each red dot marks the center of the hub region in one subject. The center of the hub was defined by the voxel with the highest weighted-sum of probabilistic streamlines from all seeded FC areas.

## Discussion

In this study we mapped out the FC inputs to different subregions within the rACC: Site 1–3 were in area 32. Site 1 was the closest to the sACC. It receives the strongest inputs from the vmPFC and the FP. Site 2 receives the strongest inputs from the FP, while site 3 receives the strongest inputs from vmPFC, vlPFC and dlPFC. Site 4 was at the conjunction of areas 32 and 24. It receives the strongest inputs from the OFC, the dlPFC, the dmPFC and the vlPFC. Sites 5 and 6 were in area 24. Site 5 receives the strongest inputs from the OFC and the vlPFC. Site 6 was the closest to the midcingulate cortex. It receives the strongest inputs from the dlPFC, the FEF and the premotor cortex. Site 4 stands out as having the highest number of input areas with the most diverse functional associations. Together with the projection strength patterns across sites, these results suggest that site 4 marks a hub region in the rACC. The dMRI tractography in the NHP demonstrated that streamlines from FC converged in a location comparable to site 4. Using the same tractography methods in human, we found that streamlines from FC also converge in a similar position in the human rACC. Thus, using a cross-species, multimodal approach we demonstrate the existence of a hub in the rACC in NHPs and, in a similar position, a likely hub human rACC.

### Species homologies

Comparative studies have utilized spatial location, cytoarchitectonics, connectivity pattern and functional activation to relate homologous areas between human and NHP. The numerical designations of FC areas in early human and NHP atlases (Brodmann, 1909; Walker, 1940) have been found most consistent in the dlPFC, while major disagreement exists in ventral and orbital areas (Barbas & Pandya, 1989; Petrides, Tomaiuolo, Yeterian, & Pandya, 2012). Such inconsistency has been further studied and alleviated based on fine-grained cytoarchitectonics and connection patterns (Carmichael & Price, 1995; Mars et al., 2018; Ongur, Ferry, & Price, 2003; Petrides & Pandya, 1994; Petrides et al., 2012). The atlases we used in this study, one by Paxinos et al. (2000) for NHP and the other by Petrides et al. (2012) for human, represent the most up-to-date knowledge of cross-species homology. These atlases were chosen to ensure best possible and accurate comparisons cross species. There are three major divisions of the ACC in NHPs (Morecraft & Tanji, 2009): the sACC (area 25), the pregenual ACC (pACC, area 32 and rostral part of area 24) and the midcingulate (caudal part of area 24). The pACC together with the rostral part of the midcingulate is also referred to as the dACC. In human neuroimaging studies, the definition is less precise. Different from the NHP terminology, the human dACC does not include parts of the pACC but only the rostral part of the midcingulate. The pACC is often referred to as the rACC. To avoid confusion, in this study we use rACC to refer to pACC in both NHPs and humans.

An important issue is whether the hub is in a homologous region within the rACC. We base our assessment on the surface features and cytoarchitecture of the ACC. The cytoarchitectonic regions of the human cingulate cortex have been found broadly consistent with the NHP region (B. A. Vogt, 2009a; Vogt, Vogt, & Laureys, 2006; Vogt, Nimchinsky, Vogt, & Hof, 1995). The two areas in this study, 32 and 24, occupy the genu of the cingulate cortex in both species. The hub region found by tract tracing was at the conjunction of NHP areas 32 and 24. In human, this conjunction region is located at the rostral-most edge of the cingulate sulcus (Vogt et al., 1995). Thus, using this surface feature as guidance, we compared the human dMRI result (Fig. 8) with a human brain atlas (Mai et al., 2008) to verify that the highest convergent connectivity was near the conjunction of areas 32 and 24.

### Translation of results between species via dMRI

Instead of a direct comparison between NHP tracing results and human diffusion tractography, we adopted diffusion tractography in NHP as an intermediate step. Comparing tracing and dMRI results in the same species ensures that dMRI captures the same pathways found by tract tracing. The direct way to replicate the tracing results was to seed at the location of each injection site and trace streamlines to the FC. However, using this method, we found disproportionally fewer streamlines exiting to cortical areas as compared to those that remained within the cingulum bundle (Fig. S2). This problem is due to the dominant fiber orientation in the cingulum bundle. Fibers are highly aligned in the cingulum bundle along the anterior-posterior axis. Consequently, the diffusion signals are strongly anisotropic towards the anterior-posterior direction, while being disproportionally weak in the other directions. Thus, streamlines from the ACC seeds have very low probability of leaving the cingulum bundle. The low number of streamlines reaching different FC areas result in low statistical power for measuring the connectivity pattern. To address this problem, we used seeds in the FC areas instead of the ACC. This way, more streamlines can be traced between an FC area and the ACC, providing sufficient statistical power for estimating the connectivity strength between each area and each ACC voxel (Fig. S2C).

### Input patterns vs. cytoarchitectonic patterns

The ACC areal divisions alone cannot fully account for the projection patterns we observed at each site. If the projections were constrained by the cytoarchitectonics of each rACC site, there would be little overlap between the input patterns to sites in area 32 and to those in area 24. Importantly, inputs to site 4 (the hub), positioned at the junction of areas 32 and 24, would equally reflect similarity to the inputs to the neighboring sites 3 & 5. However, this was not the case. First, most areas that project to area 32 also project to area 24, though with varying strength (Fig. 3). For example, comparing the sets of areas before the 100% cutoff line for sites 1 and 6 in Fig. 3, we found that areas 25, OPro, 13, 47O, OPAl, 9M, 9L, 9/32, 8A, 8B, 11 and 47L all project to both sites. Indeed, any pair of sites share common input areas. Second, the input pattern of site 4 was not a simple sum of the input patterns of its neighboring sites 3 & 5. Indeed, site 4 receives strong inputs from areas 8B, 9M, 9/32, 14O and 14M, which do not project strongly to site 3 or 5 (Fig. 5). The inputs to site 4 are uniquely diverse and involves a high number of areas from many FC regions (Figs. 3, 4, & 5). Thus, rather than being predicted by cytoarchitectonics, inputs to the rACC are better understood via three spatial patterns across all sites (Fig. 6).

### The characterization of a network hub by anatomical projection patterns

The concept of hub originates from graph theory in brain network analysis (Sporns, 2011). Theoretically, a hub is the node of the highest degree in a network, i.e. with outstandingly numerous connections with the other nodes. In modularized networks as those in the brain, the hub facilitates communication between functional modules. Neuroimaging studies have identified the rACC as a hub of the brain’s global network (Buckner et al., 2009; Hagmann et al., 2008). However, the rACC is a large region under the scope of anatomical analysis. Anatomical inputs vary across subregions within the rACC. The hub-like connectivity observed in neuroimaging studies may simply reflect the sum of connections over all of rACC’s subregions. The question is then whether a hub exists as a confined subregion within the rACC.

In this study, site 4 showed two defining features of a hub: high degree of inputs and a position in the network that facilitates cross-module integration. High degree is reflected by the high number of areas with strong projections to this site (Figs. 3–5). To demonstrate cross-module integration, we define functional modules in an empirical manner. The standard graph theory definition requires all-to-all connectivity measured between FC areas (Sporns, 2011). This is impractical with tract tracing experiments. Therefore, instead, we used functional regions of the FC to approximate functional modules. Site 4 marks the most integrative zone in the rACC where inputs from the functional regions converge. First, site 4 is in the central location of two projection gradients. One is formed by inputs from the vmPFC and the FP (Fig. 6A). These are regions associated with emotion and decision making (Joyce & Barbas, 2018; Piray, Toni, & Cools, 2016; Tsujimoto, Genovesio, & Wise, 2010). They project strongly to sites 1–3, less so to sites 4 and 5, and the least to site 6 (Fig. 6A). Inputs from the FEF and the premotor cortex are part of the second gradient. These are motor control regions that project strongly to site 6, less so to sites 4 and 5, and the least to sites 1–3 (Fig. 6B). At each end of the two gradients, the inputs are predominantly associated with emotion-or motor-related functions. Site 4 is in the intermediate transitioning zone of both gradients. Inputs are more balanced between gradients at this site. Second and importantly, a third gradient peaks at site 4. Input strength from regions associated with higher cognition gradually increases from site 1–4 and decrease from site 4–6 (Fig. 6C). The peaking inputs at site 4 allows higher cognition to maximally interface with emotion and motor control. Therefore, site 4 enables communication between all three functional modalities. Together with the high degree of inputs, the integrative nature of site 4 makes it a hub in the prefrontal network.

### Implications of FC afferent input patterns on ACC functions

There has been a dichotomy in the classical interpretation of ACC functions, such that emotional and cognitive influences affect the ventral and dorsal ACC separately (Bush, Luu, & Posner, 2000). Classically, the ventral ACC is attributed with “limbic” functions, e.g. visceral responses, emotion, and memory (Buckner, Andrews-Hanna, & Schacter, 2008; Etkin et al., 2015; Mayberg et al., 1999; Papez, 1995; Vogt, Finch, & Olson, 1992). The dorsal ACC is linked with executive control, typically for choosing from conflicting actions and monitoring behavioral outcomes (Botvinick, 2007; Holroyd & Coles, 2002; MacDonald, Cohen, Stenger, & Carter, 2000; Pardo, Pardo, Janer, & Raichle, 1990). However, this coarse dichotomy cannot entirely account for the complex activation patterns of the ACC across an increasing body of experiments. Recent theories have acknowledged the influence of reward and economic evaluation on the executive component of the dorsal ACC function (Botvinick & Braver, 2015; Kolling et al., 2016; Shenhav et al., 2016), even though reward processing is classically thought of as a ventral ACC/vmPFC function. Meanwhile, the effect of cognitive regulation over emotion in the rostral/ventral ACC has also been addressed in the literature on fear extinction (Etkin et al., 2015; Klumpp et al., 2017).

The rACC hub provides an alternative view to the ventral/dorsal dichotomy. Instead of associating each subregion with one function to support serial computation, the ACC may be better understood through the functional integration by its subregions. In neuroimaging studies, the rACC as a large region showed high degree of structural and functional connectivity with the rest of the brain (Buckner et al., 2009; Hagmann et al., 2008). In this study our results further demonstrated that the integration can be carried out in a precise subregion. Site 4 is not simply a conjunction between the ventral and dorsal ACC, but a hub that routes information across functional modules. The inputs to the hub contain strong projections from the vlPFC, the dlPFC and the dmPFC. These regions are critical for higher cognition, such as social behavior (Stalnaker, Cooch, & Schoenbaum, 2015), decision making (Kable & Levy, 2015; Sakagami & Pan, 2007; Wallis, 2007), learning (Atlas, Doll, Li, Daw, & Phelps, 2016; Schuck, Cai, Wilson, & Niv, 2016), attention (Corbetta & Shulman, 2002; Uddin, 2015) and working memory (D’Esposito & Postle, 2015; Miller & Cohen). Thus, the hub can route the outputs of higher cognitive functions to emotion and executive processing within the ventral and dorsal ACC. The hub may be uniquely positioned for evaluating and arbitrating between these processes (van den Heuvel & Sporns, 2013). The detailed mechanisms may be further investigated through connections of the hub and the other subregions of the ACC.

### Implications on the pathophysiology of psychiatric disorders

From a network perspective, most psychiatric disorders are seen as a consequence of network imbalance rather than localized deficits (Menon). In light of this view, damage to a hub region can cause disconnection between a wide range of functional modalities, and correspondingly, a spectrum of affective and cognitive disorders. The disconnection hypothesis is in line with the broadly observed rACC abnormality in various diseases, including major depression disorder (MDD) (Mayberg et al., 1997; Pizzagalli, 2011), obsessive-compulsive disorder (OCD) (Beucke et al., 2014; Tadayonnejad et al., 2017), attention deficit hyperactivity disorder (Tomasi & Volkow), and posttraumatic stress disorder (Bryant et al., 2008; Kennis, Rademaker, van Rooij, Kahn, & Geuze, 2015; Patel et al., 2012). Based on the FC areas sending convergent inputs to the hub, we propose that dysconnectivity with the hub may be key to understanding the imbalance between goal directed control, emotion and higher cognition in these disorders. MDD and OCD both show treatment response in the rACC activity (Chakrabarty, Ogrodniczuk, & Hadjipavlou, 2016; Fullana et al., 2014; Mayberg et al., 1997; O’Neill & Schultz, 2013; Pizzagalli, 2011). The distinction in their pathophysiology lies in the type of networks involved: MDD engages the networks for self-reference and cognitive control (Pizzagalli, 2011), while OCD engages those for reward-driven and goal-directed behaviors (Milad & Rauch, 2012). The hub connects a majority of FC areas involved in the above networks, which makes it a site prone to damage in both disorders. Moreover, the precise pattern of its anatomical connections provides important information for targeting disorder-specific disconnections and affected areas.

## Acknowledgements

The anatomic tracing studies and dMRI analyses were supported by NIH/NIMH grants MH106435, MH045573 and U01-MH109589, and the UK Medical Research Council grant MR/L009013/1. Collection of dMRI animal data was supported in part by the Center for Functional Neuroimaging Technologies (P41-EB015896), Shared Instrumentation Grants S10RR016811, S10RR023401, S10RR019307, and the Human Connectome Project (U01-MH093765). We thank Anna Borkowska-Belanger for expert technical support. The authors declare no conflicts of interest.

---

**Figure S1.**
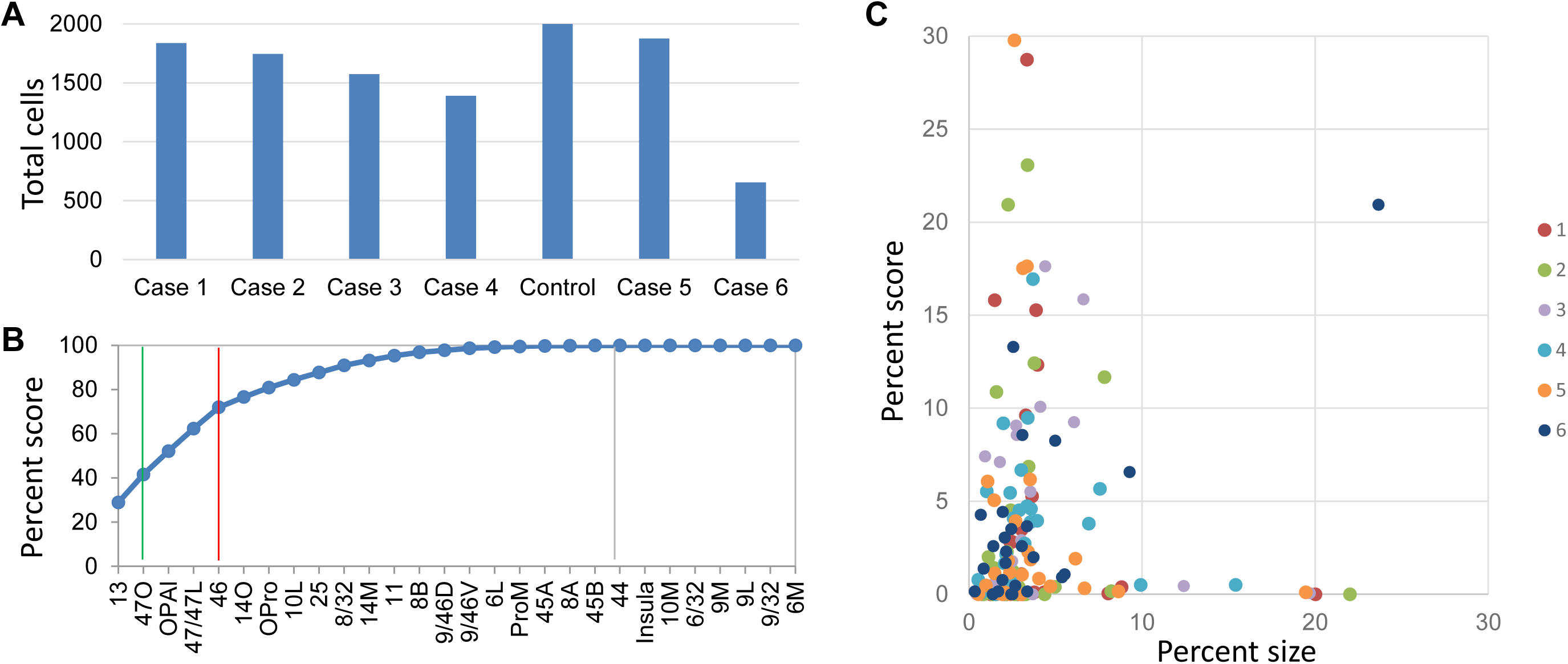
Percent scores of retrogradely labeled cells were not dominated by the size of the injection site or that of the FC areas. (A) Total number of labeled cells in each case. (B) Cumulative percent cell count across areas, showing only 2 areas contributing to 50% and 5 areas 75% of total inputs. Cutoff remarks: green line = 50%, red line = 75%, grey line = 100%. Areas after 100% are in a random order. (C) Percent scores of labeled cells projecting to each site were not determined by the size of FC areas. The size was measured as the volume of each area divided by the total volume of all areas. Colors mark different sites.

**Figure S2.**
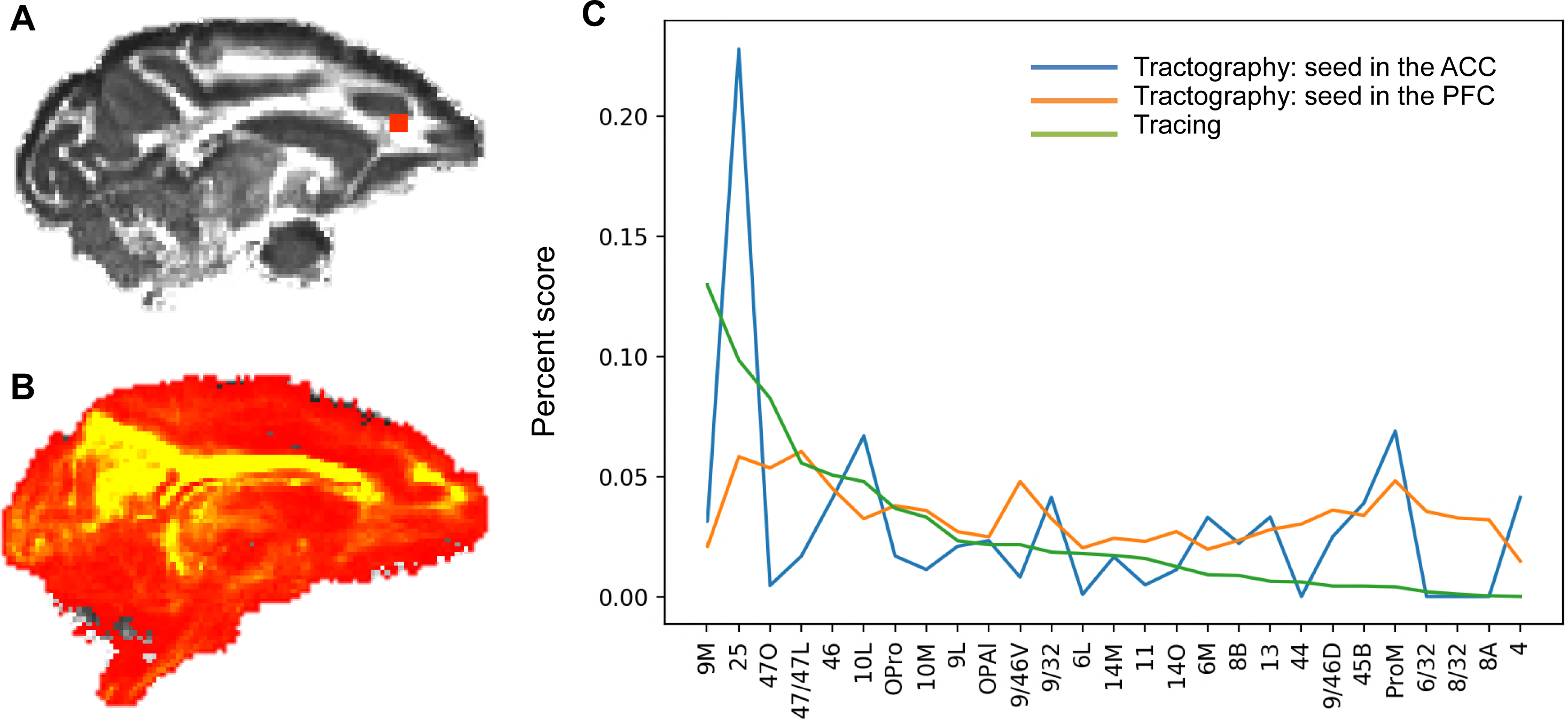
Difference in the tract strength pattern due to seeding procedures. (A) A seed mask in the rACC. (B) Probabilistic map of the streamlines from the rACC seed to all the other voxels in the brain, showing the dominant high values in the cingulum bundle. (C) Comparison of the probabilistic streamline distributions with the tract tracing result. The percent score for tracing is based on cell counts, and that for tractography is based on the tract density between the seed and the target masks (values in the fdt_path output by FSL). The tract strength by seeding the FC areas was more correlated (Pearson’s coefficient = 0.42, p < 0.05) with the tract tracing result than that by seeding the rACC (Pearson’s coefficient = 0.17, p > 0.05).

